# Mislocalisation of TDP-43 to the cytoplasm of either neurons or oligodendrocytes causes axonopathy and dysmyelination

**DOI:** 10.1101/2025.11.17.688801

**Authors:** Marcus Keatinge, Sandra Constantinou Juhasz, Toby Shaw-McGrath, Julia Albayrak, Daniel Soong, Katy Marshall-Phelps, David Story, Laura Hoodless, James Cooper, Rafael G Almeida, Tim Czopka, Giles Hardingham, David A Lyons, Siddharthan Chandran

## Abstract

Neurons with large, myelinated axons are vulnerable to degeneration across the amyotrophic lateral sclerosis (ALS) frontotemporal dementia (FTD) disease spectrum. The defining molecular pathology of this spectrum is mislocalisation of the RNA binding protein TDP-43 from the nucleus to the cytoplasm. Even though cytoplasmic TDP-43 is prevalent in neurons and oligodendrocytes, how these molecular pathologies contribute to neurodegeneration remains unclear. Here we developed humanised zebrafish in which we restricted TDP-43 to the cytoplasm of either neurons or oligodendrocytes. Cytoplasmic TDP-43 restricted to neurons led to a severe axonopathy, with extreme distal axonal swelling and reduced myelination. Axonopathy was mirrored by cell type specific loss of TDP-43 function from neurons, pointing to loss of function, rather than a toxic gain of function of mislocalised TDP-43, as driving this phenotype. Preventing oligodendrocyte differentiation and myelination when TDP-43 was mislocalised in neurons exacerbated axonopathy, indicating that oligodendrocytes limit neurodegeneration. Indeed, when TDP-43 was mislocalised to the cytoplasm of oligodendrocytes this also led to axonopathy and reduced myelination, pointing to complex contributions of neurons and oligodendrocytes to neurodegeneration.

## Introduction

Neurons in the central nervous system (CNS) are supported by oligodendrocytes that wrap myelin around axons to support action potential conduction^1^. Myelinating oligodendrocytes also maintain neuronal homeostasis by providing metabolic, antioxidant, and trophic support^2^. Experimental models that disrupt the composition of myelin or that induce damage or loss of myelin can lead to axonal damage, further highlighting the importance of myelinating oligodendrocytes to neuronal integrity^3–5^. In addition, oligodendrocytes are increasingly recognized as being disrupted in and actively regulating pathology across neurodegenerative disease (NDD)^6,7^. For example, TDP-43 proteinopathies, which include the amyotrophic lateral sclerosis/frontotemporal dementia (ALD/FTD) spectrum, represent a group of neurodegenerative diseases with significant oligodendrocyte dysfunction^7^. Neurons with large, myelinated axons are particularly vulnerable to degeneration in ALS/FTD and white matter abnormalities have been recorded in people carrying ALS/FTD-linked mutations before symptom onset^7–11^. Furthermore, associations between myelin gene variants and sporadic disease, as well as a downregulation of myelin genes, have been seen in patient tissue^12,13^. Moreover, patient-derived oligodendrocytes are toxic to healthy neurons *in vitro*, and oligodendrocyte-specific knockouts of ALS genes cause axonal damage in mice^14,15^.

The molecular hallmark of ALS/FTD is the mislocalisation of TDP-43 from the nucleus to the cytoplasm, which is prominent in both neurons and oligodendrocytes in disease^9^. Under normal conditions nuclear TDP-43 regulates RNA processing, transport, and stability^16^. In ALS/FTD, its mislocalisation to the cytoplasm could influence neurodegeneration either via loss of its core nuclear functions and/or gain of toxic functions through its ectopic localisation^16^. Despite the cytoplasmic accumulation of TDP-43 in both neurons and oligodendrocytes in disease, their relative contribution to neurodegeneration *in vivo* remains unclear. This is due, in part, the difficulty of expressing mislocalised TDP-43 at physiological levels in specific cell types and directly visualising the consequences on cellular integrity over time *in vivo.* To overcome this, we developed humanised zebrafish models, in which we mislocalised TDP-43 to the cytoplasm of either neurons or oligodendrocytes. To do so, we used a mutant form of TDP-43 (Δnls) unable to localise to the nucleus, maintaining control of physiological levels of expression by replacing zebrafish TDP-43 with the human mutated (Δnls) form. By combining this strategy for cell-type-specific humanisation with the suitability of zebrafish for live imaging neurons, axons and their associated myelin, we find that cytoplasmic mislocalisation of TDP-43 in either neurons or oligodendrocytes leads to severe axonopathy and dysmyelination. We find that neuronal cell-type specific loss of TDP-43 recapitulates the axonopathy observed upon mislocalisation of TDP-43 to the neuronal cytoplasm, indicating that this phenotype is driven by loss of function, rather than toxic gain of function. Furthermore, we find that preventing oligodendrocyte development and subsequent myelination worsens the axonopathy caused by neuronal cytoplasmic TDP-43 mislocalisation.

## Results

### Developing zebrafish models of TDP-43 proteinopathy

Zebrafish have two copies of the human TDP-43 gene (TDP-43 and TDPL) which are functionally redundant to each other^17^. Previous studies have mutated the nuclear localisation signal of the endogenous zebrafish TDP-43 using the Δnls mutation, which prevents TDP-43 from entering the nucleus, leading to permanent cytoplasmic mislocalisation^18^. Homozygous TDP-43 Δnls/Δnls leads to severe pathology but only on a TDPL null background^18^.

Importantly, wildtype human TDP-43 can functionally substitute for the zebrafish TDP-43 protein, confirming conserved function across species^17,19^. Here we aimed to assess how mislocalisation of human TDP-43 to the cytoplasm affected zebrafish neurons and oligodendrocytes. To do so we adapted the UFLIP approach^20^ to replace zebrafish TDP-43 with a full-length RFP-tagged human TDP-43 harbouring the Δnls mutation to cause cytoplasmic localisation of the human TDP-43 protein (hTDP-43). We first knocked in RFP-hTDP-43 Δnls into the endogenous zebrafish TDP-43 locus, such that zebrafish TDP-43 is exchanged for the RFP-hTDP-43 Δnls in every cell in a heritable manner (herein referred to as Δnls^Ubi^, Figure 1a and methods section for detailed explanation of line construction). Analysis of RFP in Δnls^Ubi^ showed TDP-43 to be expressed ubiquitously as expected (Figure 1b). To validate the efficiency of this approach, we performed western blots on larval heads taken at 5dpf using anti-TDP-43 antibodies which recognised both zebrafish and human TDP-43. Endogenous zebrafish TDP-43 was only detected in controls whilst RFP-Δnls TDP-43 was only detected in Δnls^Ubi/-^ confirming full humanisation of the zebrafish TDP-43 expression (Figure 1c).

**Figure 1.**
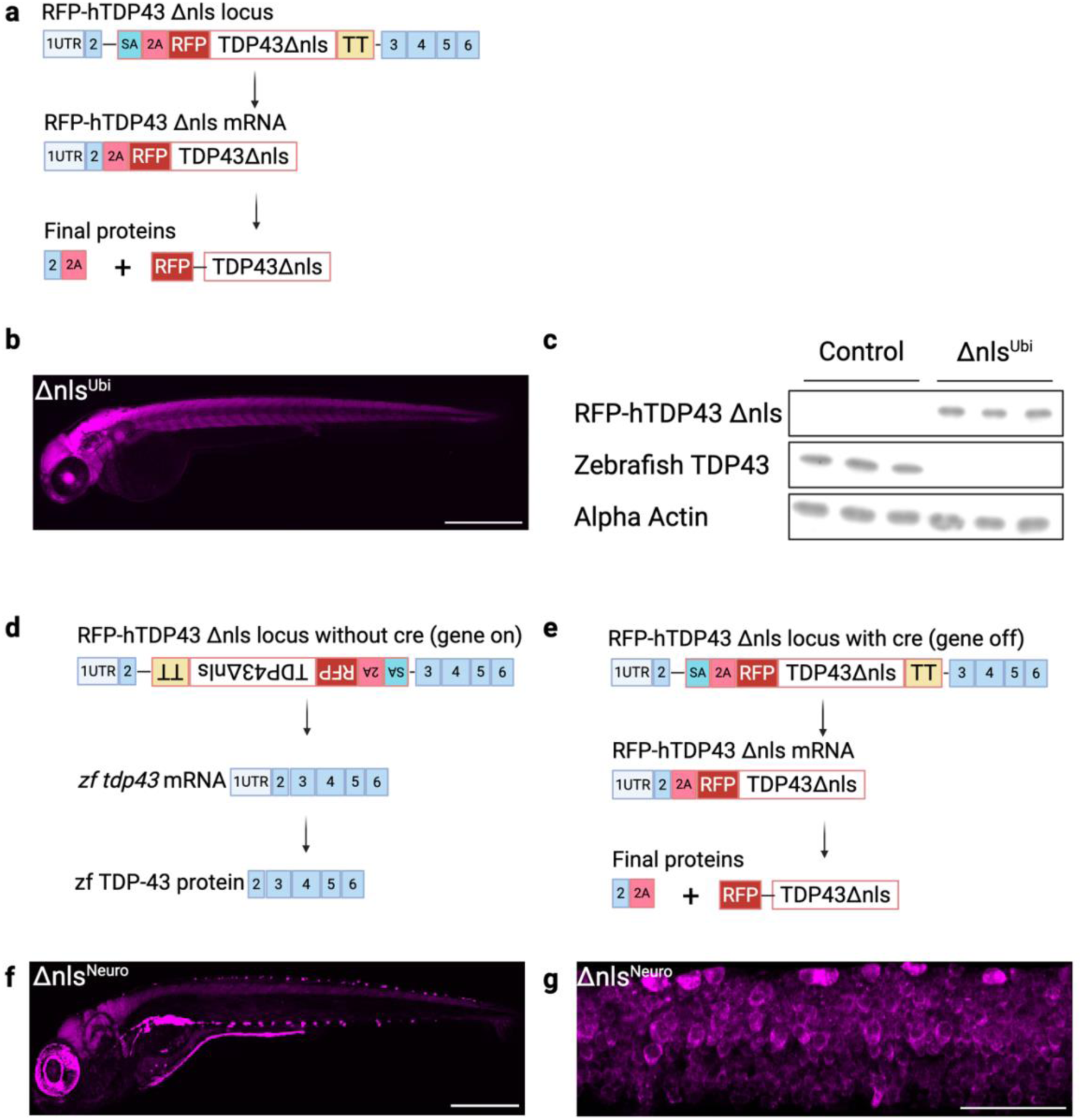
Humanising zebrafish TDP43 to express human cytoplasmic TDP43 in a physiologically relevant manner. **a.** Schematic outlining gene expression of ubiquitous RFP-hTDP43 Δnls (Δnls^Ubi^). The zebrafish UFLIP cassette has previously been knocked into intron 2 of the endogenous zebrafish TDP43 gene, in its active confirmation (gene off). The UFLIP cassette encodes a splice acceptor site (SA), a self-cleaving peptide (P2A), full length RFP fused to full length human TDP43 carrying the Δnls mutation (Δnls), followed by a super transcriptional terminator (TT). In the active (gene off confirmation) the UFLIP cassette is retained within the mRNA whilst the super transcriptional terminator prevents any further downstream transcription of endogenous exons 3-6. The novel mRNA (endogenous exon 2 and the UFLIIP cassette) is then translated, and self cleaves into endogenous TDP-43 exon 2 (which lacks function) and full-length RFP- hTDP43 Δnls. **b.** Δnls^Ubi^ individuals show RFP-hTDP43 Δnls is expressed ubiquitously at 2dpf as expected. **c.** Western blot analysis reveals full conversion of WT endogenous TDP43 to RFP- hTDP43 Δnls in the Δnls^Ubi/-^individuals. The RFP- hTDP43 Δnls band ran slightly higher than expected which has previously reported^18^. Furthermore, RFP- hTDP43 Δnls levels are reduced compared to WT control TDP-43 levels suggesting the Δnls protein is being actively cleared, as previously reported^18^. **d.** Schematic outlining gene expression of endogenous TDP-43 in control conditions “gene on” conditions. The zebrafish UFLIP cassette has previously been knocked into intron 2 of the endogenous zebrafish TDP-43 gene, in an inactive confirmation (gene on). The UFLIP cassette encodes a splice acceptor site (SA), a self-cleaving peptide (P2A), full length RFP fused to full length human TDP-43 carrying the Δnls mutation (Δnls) followed by a super transcriptional terminator (TT). In the absence of Cre, endogenous TDP-43 promoter drives the transcription of endogenous exons 1-6 whilst the UFLIP cassette is spliced out. Endogenous TDP-43 mRNA is translated into full length WT zebrafish TDP-43. **e.** In the presence of Cre (delivered via transgenic line) the UFLIP cassettes orientation is flipped over in situ exposing its splice acceptor site. The UFLIP cassette is now activated (gene off) and retained within the mRNA whilst the super transcriptional terminator prevents any further downstream transcription of endogenous exons 3-6. The novel mRNA (endogenous exon 2 and the UFLIIP cassette) is then translated, and self cleaves into endogenous TDP-43 exon 2 (which lacks function) and full-length RFP- Human TDP43 Δnls. **f.** Crossing the inactive Δnls line to a pan neuronal Cre line (Tg(Δnls; NBT: BFP T2A iCRE)) activates humanisation exclusive in neurons (Δnls^neuro^) denoted by RFP restricted to the brain and spinal cord. **g**. High magnification on the spinal cord with confocal microscopy demonstrates RFP-hTDP Δnls is fully cytoplasmic in ways previously reported of the Δnls construct^44,45^.

We next aimed to restrict RFP-hTDP-43 Δnls to the cytoplasm of neurons or oligodendrocytes *in vivo* to model both loss of nuclear function and potential gain of toxic cytoplasmic function in a cell type specific manner. Therefore, we took advantage of the fact that the UFLIP strategy can also be employed in a Cre recombinase dependent manner for cell-type specific activation. To restrict humanisation to specific cell types, we generated an inactive Δnls allele, whereby endogenous TDP-43 is expressed as normal (“gene on” Figure 1d) as the UFLIP allele is spliced out of the pre mRNA. However, the UFLIP allele can be activated (and endogenous TDP-43 is silenced- “gene off” Figure 1e) upon the delivery of Cre recombinase (through a cell type specific Cre Transgenic line for example). To test whether we could restrict humanisation to specific cell types we crossed the inactive Δnls line to a pan neuronal Cre line Tg(nbt:mtagBFP2-T2A-iCre) to activate RFP-Δnls hTDP-43 in neurons alone (herein referred to as Δnls^neuro^) (Figure 1d-e). Importantly, RFP was only detected in the brain and spinal cord (Figure 1f) and using confocal microscopy imaging, we saw that RFP-TDP-43 Δnls was localised to the cytoplasm (Figure 1g).

It is important to note that zebrafish possess two functionally redundant TDP-43 paralogs, TDP-43 (ENSDARG00000040031) and TDPL (ENSDARG00000004452)^17^. individuals with only a single functional copy of either paralog live conventional lifespans^17^. In our experimental groups, we compared non-transgenic controls with hTDP-43 Δnls mutants either in the presence (Δnls^/+^) or absence (Δnls^/−^) of a functional TDP-43/TDPL allele. This allowed us to determine which phenotypes might be due to purely toxic gain of function (Δnls^/+^) from those that might be a combination of both loss of function and toxic gain of function (Δnls^/−^).

In sum, our humanisation approach allows expression of human TDP-43 Δnls at physiologically relevant levels either in all cells or specific cell types of interest to determine cellular autonomy of phenotypes driven by cytoplasmic TDP-43 *in vivo*.

### Ubiquitous cytoplasmic TDP-43 expression causes neurodegeneration

To assess the consequences of mislocalisation of TDP-43, we first assessed the consequences of expressing RFP-hTDP-43 Δnls in all cells. We found that Δnls^Ubi/-^individuals with no functional copy of a either TDP-43 or TDPL phenocopied the previously described endogenous TDP-43 Δnls/Δnls zebrafish knock in line, in producing very little pigment (Figure 2a-b) and exhibiting lethality at larval stages, only living for ∼10 days^18^. Δnls^Ubi/+^ individuals which possessed at least one functional copy of TDP-43 (or TDPL, the zebrafish TDP-43 duplication) lived conventional life spans compared to controls as previously described^18^.

**Figure 2.**
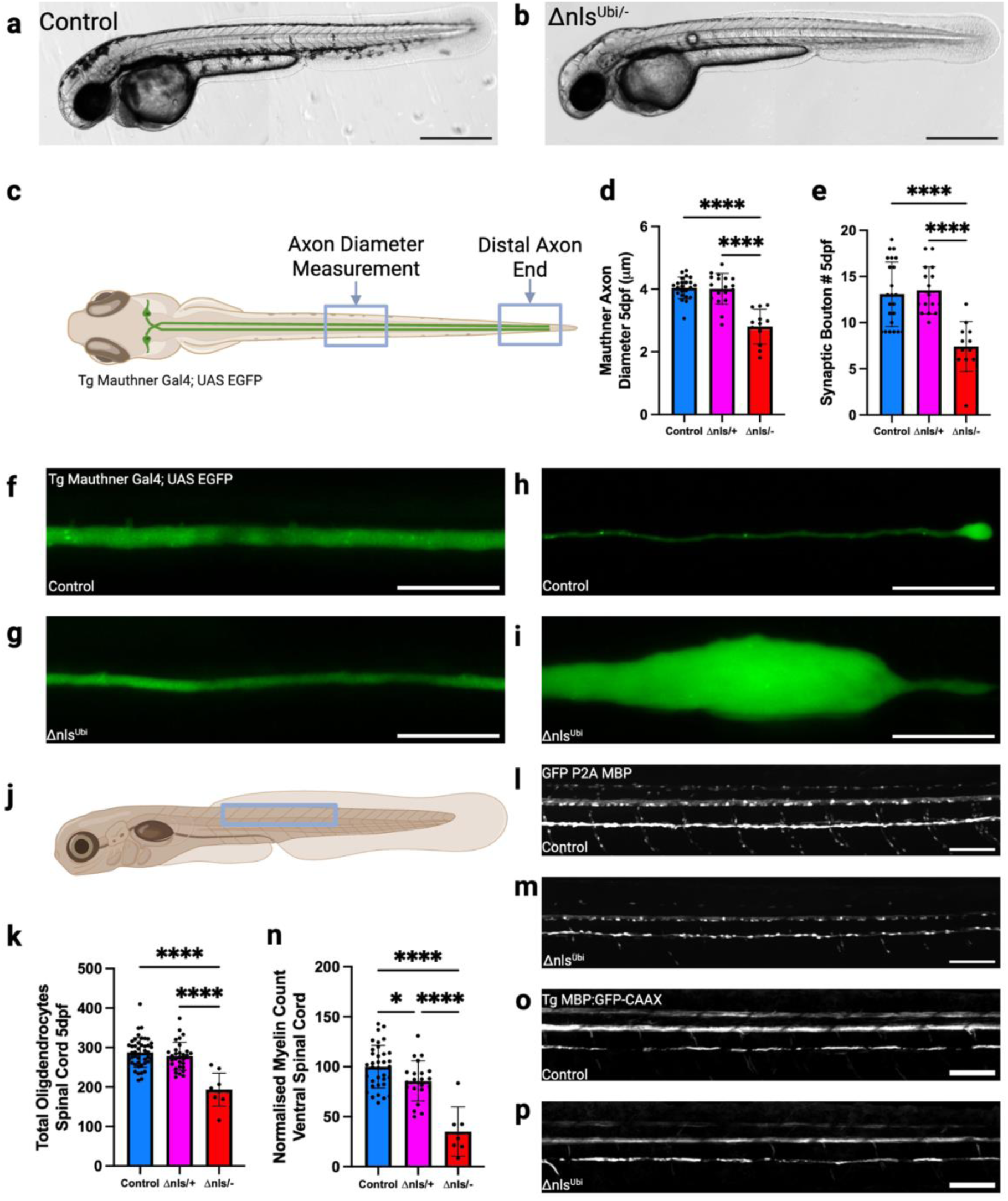
Ubiquitous cytoplasmic TDP-43 expression causes extreme axonopathy in upper myelinated motor neurons and large decreases in myelin levels. a-b. Δnls^Ubi/-^ have a lack of pigment first identifiable at 2dpf phenocopying the previously described Δnls/Δnls endogenous knock in^18^. Schematic of the Tg (Mauthner Gal4;UAS EGFP) line at 5dpf illustrating the regions under study. **c.** quantification of axon diameter of the Mauthner neuron at 5dpf in Δnls^Ubi/-^ demonstrates a 30% reduction in diameter compared to vontrol and Δnls^Ubi/+^ (p<0.0001 for both, one-way anova, n=12-23). **d.** Analysis of synaptic bouton count of the same region demonstrated a ∼55% reduction in synaptic boutons in Δnls^Ubi/-^ compared to Control and Δnls^Ubi/+^ (p<0.0001 for both, one-way anova, n=12-23). **f-g.** Representative images of the mauthner axon at the axon diameter measurement point in control (d) and Δnls^Ubi/-^ at 5dpf (scale bar=20μm). **h-i** representative images of the most distal end of the mauthner axon at 5dpf in control **(h)** and Δnls^Ubi/-^ **(i)**. The Δnls^Ubi/-^ exhibiting extreme axonopathy (scale bar=20μm). **j.** A schematic demonstrating the location of the representative images of the spinal cord. **k-m.** Representative images of oligodendrocyte reporter line KI (GFP P2A MBP, scale bar=50μm) in control **(l)** and Δnls^Ubi/-^ **(m)** spinal cord. **k**. Quantification of total oligodendrocytes numbers in the spinal cord at 5dpf. Δnls^Ubi/-^ show a ∼30% decrease in oligodendrocytes compared to Control and Δnls^Ubi/+^ (p<0.0001 for both, one-way anova, n=9-44). **n.** quantification of myelin levels in the ventral spinal cord across genotypes using Tg (MBP:GFP-CAAX). Δnls^Ubi/-^ exhibit a large decrease in myelin compared to control of 63% (p<0.0001 one-way anova, n=7-35). Of note Δnls^Ubi/+^ show a smaller yet significant decrease in myelin compared to control of 14% (p=0.0471 one-way anova, n=7-35). **n-p.** Representative images of the spinal cord of the myelin reporter TG (MBP:GFP-CAAX) from control (o) and Δnls^Ubi/-^ (p), scale bar =50μm.

In patients of the ALS/FTD spectrum, TDP-43 pathology is primarily found in neurons and oligodendrocytes^21^. Thus, we initially focussed on assessing the effects of ubiquitous cytoplasmic TDP-43 expression on these cell types.

To begin to determine the impact of ubiquitous cytoplasmic TDP-43 on neuronal health we used the transgenic reporter line Tg(Mauthner Gal4; UAS EGFP). This transgenic line labels the Mauthner neurons, a pair of upper motor neurons localised in the hindbrain that project large calibre axons through the spinal cord to drive escape responses. Importantly these neurons have attributes common with the fast fatigable central motor neurons most at risk in ALS (Figure 2c)^22,23^. Due to the transparency of larval zebrafish, this transgenic line allows us to visualise the neuron and its entire axon *in vivo* over time, in a non-invasive manner.

To assess how RFP-hTDP-43 Δnls in all cells affected neuronal morphology, we first measured the diameter of the Mauthner axons and quantified synaptic bouton number. We found that axon diameter was reduced by 30% in Δnls^Ubi/-^ compared to Δnls^Ubi/+^ and controls when analysed at 5 days post-fertilisation (dpf) and at a position mid-way along the length of the axon (circa somite 15) (Figure 2d-g). We saw that the number of pre-synaptic boutons of the Mauthner axon were decreased by ∼55% in Δnls^Ubi/-^compared to Δnls^Ubi/+^ and controls at 5dpf (Figure 2e). In addition to these phenotypes, inspection of the most distal end of the Mauthner axon at 5dpf revealed an extreme axonopathy in the form of gigantic swellings in the Δnls^Ubi/-^ (Figure 2h-i). Axons exhibiting such swellings were often retracted by several somite lengths compared to control individuals, suggesting die-back of axons, although this was variable. Over the next several days the Mauthner axons in the Δnls^Ubi/-^ underwent blebbing and frank degeneration. Of note, no significant differences in Mauthner axon integrity (diameter, synapse number or distal axon morphology) were detected between Δnls^Ubi/+^ and controls, suggesting that the mislocalisation of human TDP-43 alone is not sufficient to drive these cellular pathologies.

As myelinated axons are at risk of degeneration in ALS/FTD, we next wanted to determine how oligodendrocytes and myelin levels were affected by ubiquitous cytoplasmic TDP-43 expression. Using a newly-generated fluorescent reporter line, KI(GFP-P2A-MBP), we quantified all myelinating oligodendrocytes in the spinal cord across genotypes (Figure 2j-m). Oligodendrocyte number was decreased by ∼30% in the Δnls^Ubi/-^ compared to Δnls^Ubi/+^ and controls (Figure 2k). Subsequently we used Tg(mbp:EGFP-CAAX) to measure myelin levels, which we found were reduced by ∼60% in the CNS of the Δnls^Ubi/-^ respectively, compared to controls (Figure 2n-p). Of note, the myelin levels in Δnls^Ubi/+^ were also decreased compared to controls by 14%.

Taken together, our results demonstrate that ubiquitous expression of cytoplasmic TDP-43 at physiologically relevant expression levels reduces oligodendrocyte and myelin levels.

### Neuronal restricted cytoplasmic TDP-43 causes neurodegeneration and a reduction in myelin

TDP-43 pathology is present in both neurons and oligodendrocytes in ALS/FTD. As we had established that ubiquitous cytoplasmic TDP-43 expression led to both neuronal and oligodendrocyte dysfunction *in vivo*, we sought to determine which phenotypes were being driven by cytoplasmic localisation of TDP-43 in neurons. Therefore, we crossed our inactive (“gene on”) Δnls line to a pan neuronal Cre line that we generated, Tg (nbt:mtagBFP2-T2A-iCre) to restrict cytoplasmic TDP-43 purely to neurons (Methods).

Initial inspection of larvae showed the Δnls^neuro/-^ line did not exhibit the obvious pigment defects detected in the Δnls^Ubi/-^ and were indistinguishable from controls. This suggests that gross morphological deficits in the Δnls^Ubi/-^ are likely driven from dysfunction from different or additional cell types.

We first sought to investigate whether neuronal defects seen in the Δnls^Ubi/-^ such as reduced axon diameter and extreme axonopathy were seen when TDP-43 was restricted to neurons only. As Δnls^neuro/-^ larvae could not be sorted by visual inspection, all embryos were sorted for expression of the RFP tag, and Δnls^neuro/+^ were used as controls in experiments, given that Δnls^Ubi/+^ showed no phenotype compared to non-transgenic controls. Initial analysis of the Mauthner axon diameter at 5dpf revealed a non-significant decrease in the Δnls^neuro/-^ of 11% compared to the control Δnls^neuro/+^ (Figure 3a). We hypothesised that neuronal phenotypes may need further time to manifest, due to the Cre-dependence of neuron-specific TDP-43 mislocalisation. Therefore, we undertook the same analysis at 7dpf, which revealed a significant decrease of 27% in Mauthner axonal diameter in the Δnls^neuro/-^ compared to the control

**Figure 3.**
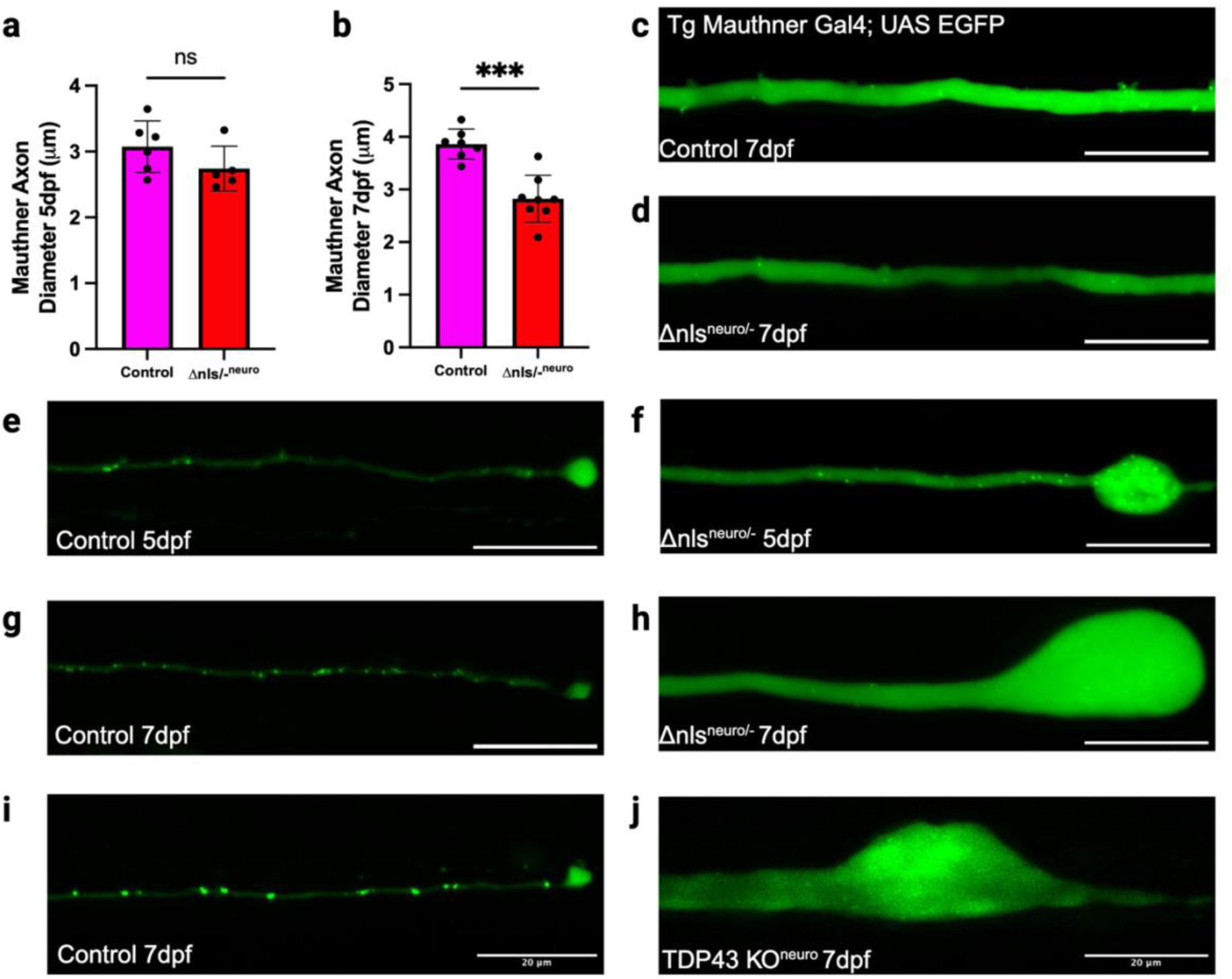
Neuronal restricted cytoplasmic TDP-43 results in neurodegeneration. **a.** Mauthner axon diameter is not significantly different at 5dpf between Δnls^neuro/-^ and control (Δnls^neuro/+^) (p=0.1708 unpaired t test, n=5-6). **b.** However at 7dpf axon diameter is reduced by 27% in Δnls^neuro/-^ compared control (p=0.0002 unpaired t test, n=8-9). **c-d**. Representative images of the mauthner axon at the axon diameter measurement point in control **(c)** and Δnls^neuro/-^ **(d)** at 7dpf (scale bar=20μm). **e-f.** Representative images of the most distal end of the mauthner axon at 5dpf in control **(e)** and Δnls^neuro/-^ **(f)**. The Δnls^neuro/-^ exhibiting partial axonopathy at 5dpf (scale bar=20μm). **g-h.** Representative images of the most distal end of the mauthner axon at 7dpf in control **(g)** and Δnls^neuro/-^ **(h)**. The Δnls^neuro/-^ exhibiting extreme axonopathy phenocopying the 5dpf Δnls^Ubi/-^ timepoint (scale bar=20μm). **i-j** Control Mauthner axon at 7dpf **(i).** Neuronal TDP43 specific KO Mauthner axon at 7dpf demonstrating dramatic swelling in like with Δnls^neuro^ at 7dpf **(j).**

Δnls^neuro/+^ (Figure 3b-d). This is a similar decrease in axon diameter to the 5dpf Δnls^Ubi/-^(Figure 2d). Analysis of the distal end of the Mauthner axon revealed axonopathy at 5dpf in the Δnls^neuro/-^ compared to control, although these were reduced in size compared to swellings exhibited by Δnls^Ubi/-^ at the same time point (Figure 3e-f). However, at 7dpf the distal end of the Mauthner axon in Δnls^neuro/-^ had swollen to similar sizes exhibited by the 5dpf Δnls^Ubi/-^ (Figure 3g-h), with each Mauthner axon retracted by several somites’ length in the Δnls^neuro/-^ compared to controls, again suggestive of axonal die-back. Our data demonstrate that extreme axonopathy exhibited in the Δnls^Ubi^ line is also prevalent in the Δnls^neuro^ line (albeit delayed).

Cytoplasmic TDP-43 is believed to cause neurodegeneration through a combination of loss of TDP-43 function and toxic gain of function, but how each mechanism contributes to neuronal loss is disputed. The absence of neuronal phenotypes in the Δnls^Ubi/+^ and Δnls^Neuro/+^ individuals, which model purely toxic gain of TDP-43 function, suggested the Mauthner neuron phenotypes of Δnls^Ubi/-^ and Δnls^Neuro/-^ are driven primarily by loss of TDP-43 function. To test this more directly, we also generated a TDP-43 cell type specific knock out line TDP-43^UFLIP KO^ (see methods for a detailed description of line generation) using the UFLIP strategy, which we crossed to our pan neuronal Cre line to KO TDP-43 specifically from neurons. The Mauthner axon swelled to gigantic proportions by 7dpf in the TDP-43^UFLIP KO; neuro/-^ line, in a similar fashion to the Δnls^Neuro/-,^ confirming the TDP-43 loss of function is the primary driver of cytoplasmic TDP-43 dysfunction (Figure 3i-h)

We then sought to investigate whether neuronally restricted cytoplasmic TDP-43 could modulate oligodendrocytes in a non-cell autonomous manner. In contrast to the Δnls^Ubi/-^ which had a ∼30% reduction in oligodendrocytes (Figure 2k), restricting cytoplasmic TDP-43 to neurons using the Δnls^neuro/-^ line did not modulate oligodendrocyte numbers compared to controls at 5dpf (Figure 4a). However, despite possessing conventional numbers of oligodendrocytes at this stage, restricting TDP-43 pathology to neurons did lead to a ∼30% decrease in myelin in the CNS of the Δnls^neuro/-^ compared to control, assessed using the transgenic reporter Tg(mbp:EGFP-CAAX) (Figure 4b-d).

**Figure 4.**
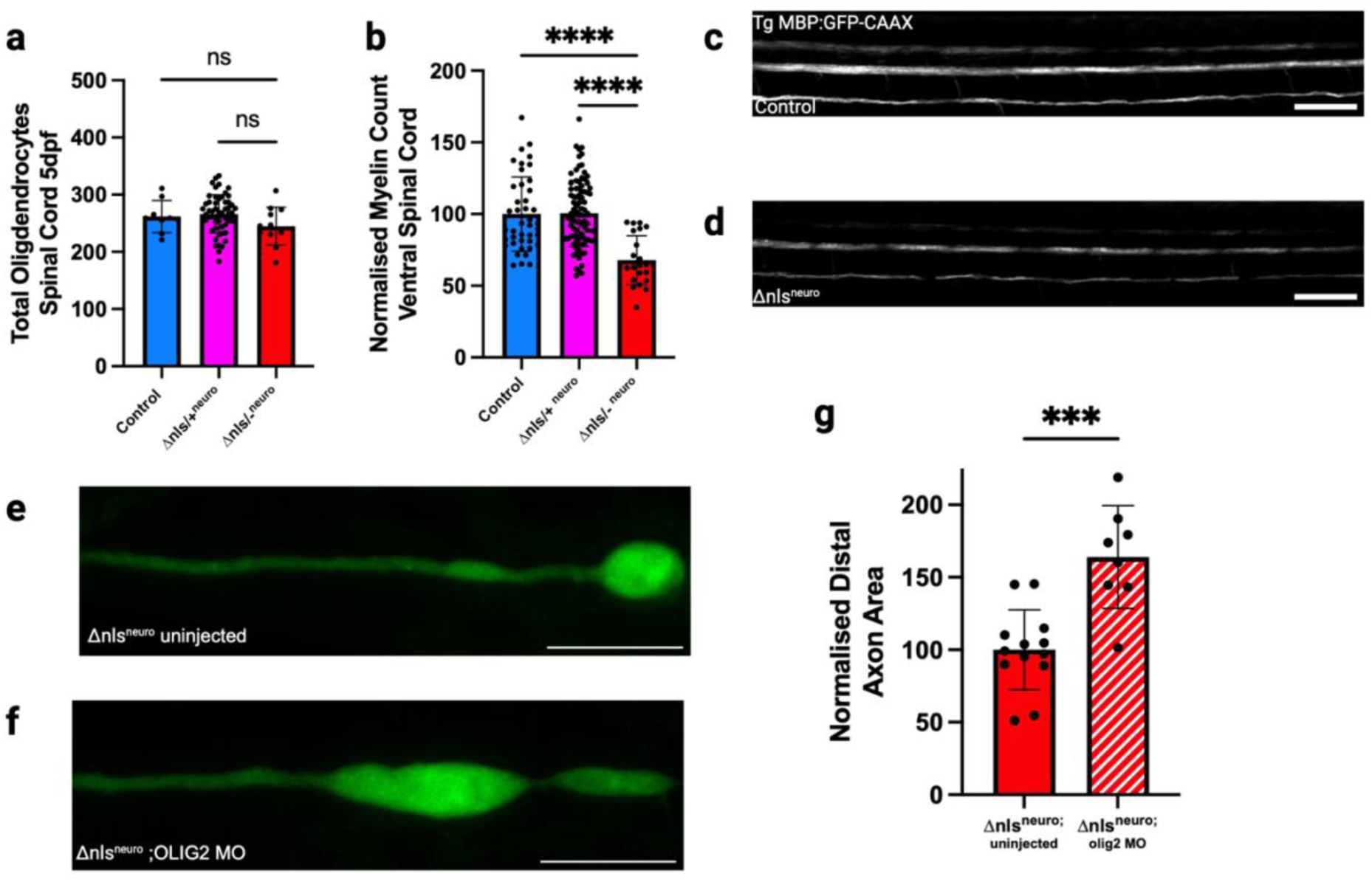
Myelin is reduced but still neuroprotective when cytoplasmic TDP-43 is restricted to neurons. **a.** Quantification of total oligodendrocytes numbers in the spinal cord at 5dpf. Oligodendrocyte numbers are not significantly different between genotypes at 5dpf (p=0.4865, one-way anova, n=9-53) **b.** Quantification of myelin levels in the ventral spinal cord across genotypes using Tg (MBP:GFP-CAAX). Δnls^neuro/-^ exhibit a decrease in myelin compared to controls and Δnls^neuro/+^ of 32% (p<0.0001 one-way anova, n=23-88). **c-d.** Representative images of the spinal cord of the myelin reporter TG (MBP:GFP-CAAX) from control **(c)** and Δnls^neuro/-^ **(d),** scale bar =50μm. **e-f.** Representative images of the distal end of the mauthner axon at 5dpf in uninjected Δnls^neuro/-^ **(e)** and Δnls^neuro/-^ ;OLIG2 KD **(f)**, scale bar= 20μm. **g.** Quantification of mauthner axon swellings at 5dpf in uninjected Δnls^neuro/-^ and Δnls^neuro/-^ ;OLIG2 KD. Preventing myelination in Δnls^neuro/-^ increased axon swellings by 64% compared to uninjected Δnls^neuro/-^ (p=0.0002 unpaired t test with welchs correction, n=8-13).

In our neuronal model, this reduction in myelin precedes the extreme axonopathy of the Mauthner axon. As myelin deficits are known to lead to axonal damage in experimental model systems, we reasoned that a reduction in myelin might be contributing to the subsequent axonopathy in the context of neuronal cytoplasmic TDP-43. Alternatively, neuronal cytoplasmic TDP-43 might non-autonomously disrupt oligodendrocyte integrity such that they actively damage axons, whereby their removal might relieve neurodegeneration. To begin to disentangle between these possibilities, we prevented myelination in the neuronal restricted Δnls TDP-43 individuals by knocking down Olig2^24^ as previously described. Preventing myelination exacerbated the distal axonopathy of Mauthner axon increasing the size of swellings by 64% compared to controls (Figure 4e-g). Taken together, our data suggests that the myelin that remains present in the context of neuronal cytoplasmic TDP-43 helps protect axons from more severe axonopathy.

### Oligodendrocyte restricted cytoplasmic TDP-43 reduces myelin levels and perturbs axons

In the ALS/FTD spectrum, cytoplasmic TDP-43 mislocalisation is also detected in oligodendrocytes whilst myelinated axons are sensitive to degeneration^8,21^. Therefore, we reasoned that restricting cytoplasmic TDP-43 to oligodendrocytes might lead to both myelin and neuronal dysfunction.

We restricted cytoplasmic TDP-43 to oligodendrocytes (herein referred to as Δnls^oligo^) by driving Cre in the oligodendrocyte lineage using a newly generated transgenic line Tg(Olig1: iCre). We first measured the number of oligodendrocytes using KI(GFP-P2A-MBP) and found oligodendrocyte number to be unchanged compared to controls in Δnls^oligo/-^ (Figure 5a). However, CNS myelin was decreased in the Δnls^oligo/-^ by 39% (Figure 5b-d). Myelin levels CNS were not significantly different in Δnls^oligo/+^ individuals which have cytoplasmic TDP-43 in oligodendrocytes but retain a functional copy of zebrafish TDP-43, compared to controls. This further suggests that the primary mechanism leading to phenotypes at these stages is a loss of TDP-43 function, rather than a toxic gain of function due to mislocalised TDP-43.

**Figure 5.**
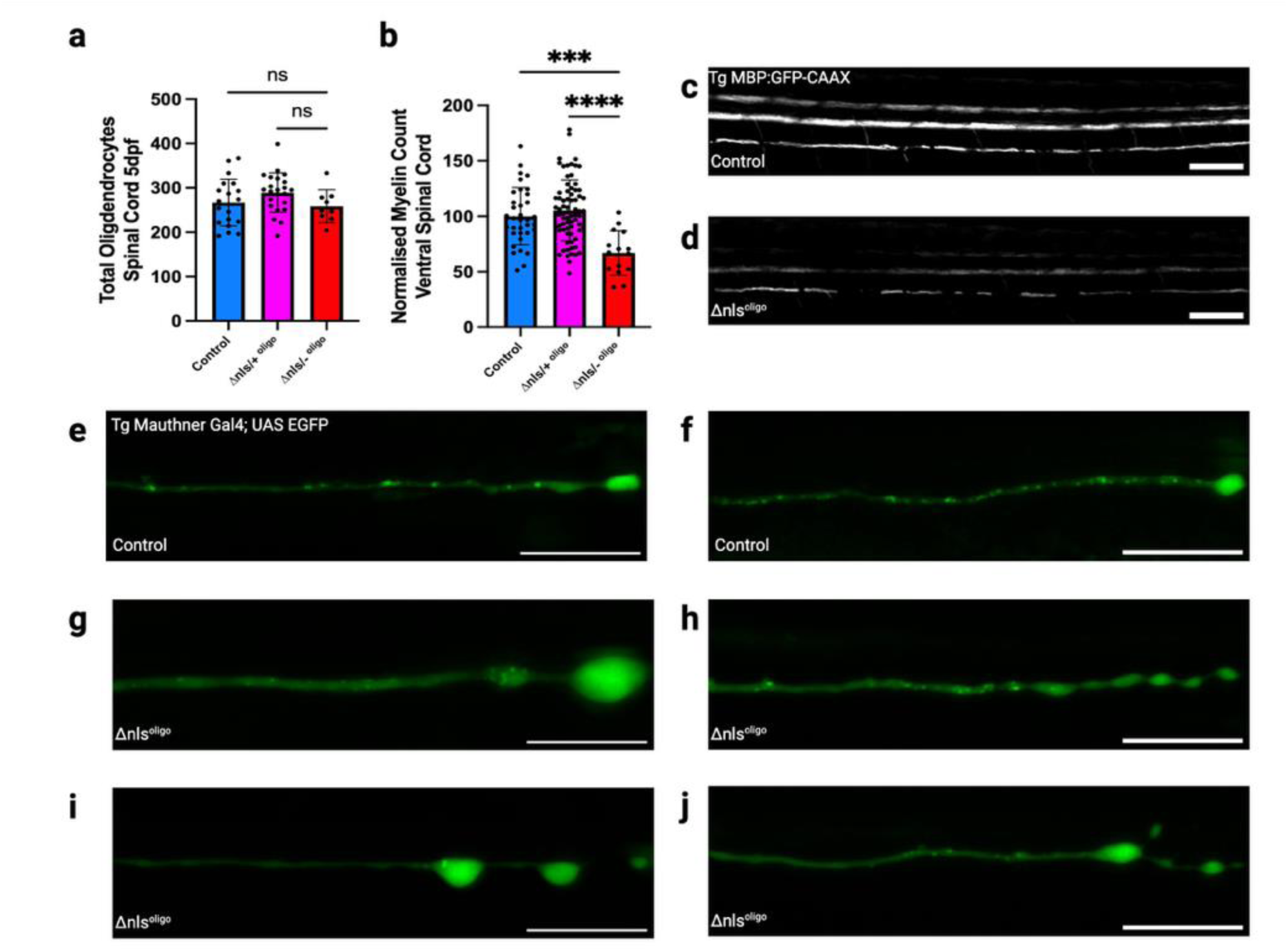
Oligodendrocyte restricted cytoplasmic TDP-43 results in axonal and oligodendrocyte dysfunction. **a.** Quantification of total oligodendrocytes numbers in the spinal cord at 5dpf in Δnls^oligo/-^. Oligodendrocyte numbers are not significantly different between genotypes at 5dpf (p=0.8944, one-way anova, n=9-23) **b.** Quantification of myelin levels in the ventral spinal cord across genotypes using Tg (MBP:GFP-CAAX). Δnls^oligo/-^ exhibit a decrease in myelin compared to controls and Δnls^oligo/+^ of 33% (p=0.0002 one-way anova, n=15-73). **c/d.** Representative images of the spinal cord of the myelin reporter TG (MBP:GFP-CAAX) from control (c) and Δnls^oligo/-^ (d), scale bar =50μm. **e-j**. Representative images of the most distal end of the mauthner axon at 5dpf in control **(e-f)** and Δnls^oligo/-^ **(g-j)** (scale bar=20μm). Δnls^oligo/-^ exhibit axonal dysfunction to varying levels compared to controls at 5dpf. Either a single swelling of a similar size to the 5dpf Δnls^neuro/-^ **(g),** multiple smaller swellings forming a beaded appearance **(h-i),** or terminated the axon at the yolk extension producing multiple sprouts **(j).**

The Mauthner neuron is highly myelinated, and our Δnls^oligo/-^ showed a reduction in myelin levels. To assess how oligodendrocyte restricted cytoplasmic TDP-43 affected axons, we examined the distal end of the Mauthner neuron at 5dpf to assess whether it exhibited any disruption to axonal morphology. The Δnls^oligo/-^ exhibited variable types of structural defects at the end of the distal end (Figure 5e-j). The Δnls^oligo/-^ distal end developed either a single swelling of a similar size to the 5dpf Δnls^neuro/-^ (Figure 5g), multiple smaller swellings forming a beaded appearance (Figure 5h-i), or terminated the axon at the yolk extension producing multiple sprouts (Figure 5j). These observations indicate that mislocalisation of TDP-43 to oligodendrocytes leads to non-cell autonomous disruption to axonal integrity.

When taken together our results demonstrated oligodendrocyte restricted cytoplasmic TDP-43 results in cell autonomous (myelin reduction) and crucially, non-cell autonomous phenotype (axon dysfunction).

## Discussion

The contributions of different cell types in driving neurodegeneration due to cytoplasmic TDP-43 are unclear, in part due to the lack of *in vivo* models restricting cytoplasmic TDP-43 to specific cell types^18,25,26^. We used our conditional model, to selectively restrict cytoplasmic TDP-43 to neurons and oligodendrocytes to investigate cellular autonomy in TDP-43 proteinopathies. As our platform leverages the endogenous zebrafish TDP-43 gene promoter this ensures expression at physiologically relevant levels. TDP-43 pathology is common in both neurons and oligodendrocytes, and oligodendrocyte dysfunction is prevalent in ALS/FTD^7^. However, the majority of *in vivo* models modulate TDP-43 exclusively in neurons, therefore the glial contributions to neurodegeneration are unknown^25^. Our findings reveal that both neuronal and oligodendrocyte-specific expression of cytoplasmic TDP-43 leads to axonal dysfunction and a reduction in myelin levels.

Of note, the large axonal swellings which develop in the Δnls^neuro^ line are similar to those previously reported in zebrafish axonal transport mutants, specifically that disrupt dynein/dynactin and retrograde transport ^27^. Indeed, TDP-43 deficiency *in vitro* in a variety of different cell lines has also been reported to result in a decrease in dynactin levels, a key axonal transport protein^28^. Furthermore, myelin dysfunction can result in axonal transport defects^29^. It is therefore plausible in the Δnls^neuro^, cargo is trafficked to the distal end of the axon, but then progressively accumulates there due to retrograde transport defects.

We also observed disruption to myelin and axons upon mislocalisation of TDP-43 to the oligodendrocyte cytoplasm. In terms of this Δnls^oligo^ context, TDP-43 deficient oligodendrocytes from mice have been reported to exhibit metabolic dysfunction in vivo^30^. Oligodendrocyte metabolic dysfunction could therefore lead to a decrease in metabolic support to axons and thus contribute to degeneration^30,31^. This could be particularly critical to large, myelinated axons with a high metabolic demand. Future efforts will be dedicated to assessing disruption to both axonal transport and metabolism in the Δnls lines. For example, zebrafish transgenic reporters of axonal transport (such as for lysosomes and mitochondria) and ATP will be developed as part of future work^32^.

Of note, both the Δnls^neuro^ and Δnls^oligo^ lead to a decrease in myelin levels and in the case of the Δnls^neuro^, the reduction of myelin precedes the giant axonal swellings. When we prevented myelination from occurring (via Olig2 KD) in the Δnls^neuro^ line, the axonal swellings were exacerbated. This suggests that myelin, even though reduced in the context of Δnls^neuro^, is still providing a neuroprotective effect. There is the potential that raising myelin levels back to those of control individuals may therefore rescue axonal phenotypes, e.g. by pharmacological targeting.

The ability of cytoplasmic TDP-43 in either neurons or oligodendrocytes to induce dysfunction in the opposing cell type in a non-cell-autonomous manner highlights the close connection between neurons and oligodendrocytes. Furthermore, it also suggests a positive feedback loop in which TDP-43 pathology propagates neurodegeneration through reciprocal neuron-oligodendrocyte interactions.

Of note, the Δnls^Ubi/+^ line which expresses the cytoplasmic human TDP-43 in all cells, whilst retaining endogenous zebrafish TDP-43 function, in essence modelling toxic gain of function, displayed minimal phenotypes and lived conventional lifespans. Moreover, our neuronal specific TDP-43 KO line phenocopied the extreme axonal swellings exhibited by the Δnls^neuro/-^. This data argues that loss of TDP-43 function is the dominant mechanisms driving neurodegeneration *in vivo*.

The previously described endogenous Δnls TDP-43 knock in zebrafish line develops cytoplasmic TDP-43 mislocalisation in every cell of the zebrafish under the control of the endogenous TDP-43 promoter^18^. The Δnls/Δnls homozygotes develop neurodegeneration, pigment defects and early lethality. However, this only occurs when on a background devoid of any function TDP-43 or TDPL. This matches the phenotypes produced by the Δnls^Ubi/-^ line, validating our humanisation approach. However using our conditional humanisation system, we were then able to deconstruct the cellular autonomy of neurons and oligodendrocytes, demonstrating the complicated role each plays in neurodegeneration, instead of oligodendrocytes merely being passive bystanders.

Finally, our humanised TDP-43 zebrafish line is versatile and not limited to just analyses of oligodendrocytes and neurons. Using different Cre lines, cytoplasmic TDP-43 mislocalisation could be restricted to endothelial cells or astrocytes for example. Cell types which both develop TDP-43 pathology in NDD^16^. Furthermore, the modified UFLIP strategy itself could be applied to other neurodegeneration-linked genes which require conditional physiological expression of toxic proteins in a cell type restricted manner *in vivo*, such as mutant alpha synuclein or tau. This would be especially useful for deconstructing the contributions of neurons and glial cells across different neurodegenerative disease models.

## Materials and Methods

### Zebrafish Husbandry

Adult zebrafish (*Danio rerio*) were housed and maintained in accordance with standard procedures in the Queen’s Medical Research Institute zebrafish facility, University of Edinburgh. All experiments were performed in compliance with the UK Home Office, according to its regulations under project licences 70/8436, PP5258250 and PP3290955. Adult zebrafish were subject to a 14/10 h light/dark cycle. Embryos were produced by pairwise matings and were staged according to days post-fertilisation (dpf). Embryos were raised at 28.5°C, with 50 embryos (or less) per 10 cm petri dish in ∼45 ml of 10 mM HEPES-buffered E3 embryo medium or conditioned aquarium water with methylene blue.

### Genome editing

TracrRNA /CrRNA (both Merck/Sigma), Cas9 protein (NEB) and injection mix made up as previously described^20,33–35^. Briefly, TracrRNA and CrRNA were combined and heated at 95°C for 5 minutes the cooled. Cas9 protein, dye and donor DNA (if required) were added then the mixture incubated at 37°C for 5 minutes before injection into the cell of the embryo at the one cell stage ^36^.

### Zebrafish line generation

#### Standard loss of function mutants

Loss of function mutants for TDP-43 (10bp del exon 3) and TDPL (56bp del +18bp ins exon 1) were generated by CRISPR-Cas9-based gene editing, using CrRNAs targeting sequence 5’ TGGGGAAGTCATCATGGTGC (TDP-43) and 5’ CATGAGAGGGGTCCGACTAG (TDPL) respectively. Each embryo was genotyped post hoc after data analysis using TDP-43 primers F 5’ GCGACCAAGATCAAGAGAGG and 5’ R ACCACGGGTGTATGGGTTAT followed by an msl1 digest. TDPL was genotyped using primers 5’ F TCGGTGTAATCATGACGGAGT and 5’ AATAAGCAGGTCGAACCCATT and genotyped by band shift on a gel due to the large size of the indel.

#### TDP-43 cell type specific KO/Δnls line generation

The TDP-43 cell type specific KO line and Δnls humanisation lines were generated with the UFLIP strategy^20^. Each UFLIP construct was made using geneweld^37^ and targeted to intron 2 of TDP-43 using the target sequence CrRNA 5’ AGAACCACTGGCCAGATTGG. The RFP-hTDP-43 Δnls UFLIP construct for humanisation was generated using a gblock (IDT) encoding cDNA for RFP-human TDP-43 carrying the Δnls mutation. This was subcloned into conventional UFLIP plasmids using restriction enzymes AatII (NEB) and SpeI (NEB).

Each UFLIP construct was assembled in either the “gene off position” or “gene on position” (Figure 1d and e). When in the “gene off position” endogenous TDP-43 expression is permanently (in a stable and heritable fashion) exchanged for expression of RFP, or RFP fused to human TDP-43 carrying the Δnls mutation (depending on UFLIP line). When in the “gene on position”, endogenous TDP-43 is expressed as normal until the addition of Cre, whereby the UFLIP construct is activated, inhibiting expression of the endogenous TDP-43, exchanging it for expression of either RFP alone (for cell type specific KO) or for expression of RFP fused to human TDP-43 carrying the Δnls mutation (for cell type specific mislocalisation to the cytoplasm). The TDP-43 cell type specific KO line was generated only in the “gene on” form whilst the RFP-hTDP43 Δnls lines were generated in both the “gene on” form (for cell type specific mislocalisation) and the “gene off” form (to create the permanent and ubiquitous TDP-43 Δnls expression for the Δnls Ubi line).

To assemble the UFLIP constructs in its “gene on” position, UFLIP-2ARFP_ HE1A-1 was combined with annealed primers using homology arms upstream A 5’cggaaaagtgaaaattttggcagggcaagtaaaagtcagaaccactggccagat, upstream B 5’aagatctggccagtggttctgacttttacttgccctgccaaaattttcactttt, downstream A 5’gaagtgggggccagtagaaaaaatccttagcgttgaaccctaatacagtaaaggg and downstream B 5’gcggccctttactgtattagggttcaacgctaaggattttttctactggccccca.

To assemble the UFLIP construct in its “gene off” position UFLIP-2ARFP_ HE1A-1 was combined with annealed primers using homology arms upstream A 5’cggaaaagtgaaaattttggcagggcaagtaaaagtcagaaccactggccagat, upstream B 5’aagatctggccagtggttctgacttttacttgccctgccaaaattttcactttt, downstream A 5’gaagtgggggccagtagaaaaaatccttagcgttgaaccctaatacagtaaaggg and downstream B 5’gcggccctttactgtattagggttcaacgctaaggattttttctactggccccca.

Plasmids where sequenced to confirm identify then injected into the cell at the one cell stage with Cas9 (NEB), the universal guide (CrRNA spacer 5’ GGGAGGCGTTCGGGCCACAG) and the CrRNA targeting intron 2 of zebrafish TDP-43 (5’ AGAACCACTGGCCAGATTGG). Injected fish carrying the blue hatching pec marker only (gene on position) or blue eyes/ubiquitous RFP expression (gene off position) were raised. Once at adulthood, injected fish were crossed to WT, and F1 individuals positive for the blue hatching pec marker (gene on) or blue eye maker/ubiquitous RFP expression (gene off) were analysed by junction PCR at both the 5’ and 3’ junction to confirm on target integration of UFLIP RFP/RFP-TDP-43 Δnls. For the UFLIP RFP- 5’ junction F 5’ GTTATTAACTTCTTTTTTTGGTGGGGCC and R 5’ GCAAACGGCCTTAACTTTCC, for 3’ junction F 5’ CCTTGGTCACCTTCAGCTTG and R 5’ TAGTTCCTTTATTTAAAAGCTCAGATTTG. For RFP-hTDP-43 Δnls gene off, 5’ junction F 5’ TAATTTAAGCCTATAATATATATATTTTC and R 5’ CCTTGGTCACCTTCAGCTTG, 3’ junction F 5’ GCAAACGGCCTTAACTTTCC and R 5’ AGGGTAATATATTCATTCAACTTCACA. For RFP-hTDP-43 Δnls gene on, 5’ junction F 5’ CAAATGTATTAACATCACAGTTGGA and 3’ GCAAACGGCCTTAACTTTCC, 3’junction F 5’ CCTTGGTCACCTTCAGCTTG and R 5’ AGGGTAATATATTCATTCAACTTCACA. The locus of each was then sequenced to validate integration with the same primers.

#### Western blot

Inhibition of endogenous TDP-43 and subsequent exchange of expression for human TDP-43 Δnls was confirmed using Western blot. Heads of 5dpf larvae where flash frozen in E3 until X4 laemlli buffer was added and the mixture heated for 5 minutes at 95°C, vortexed twice then heated for another 5 minutes at 95°C. Protein samples were run on a Novex precast gel (Invitrogen) for 90 minutes and probed with either anti TDP-43 antibodies (A260) Antibody #3449, cell signalling) that recognised both zebrafish and human TDP-43, or alpha tubulin (12G10 anti-alpha-tubulin, DSHB).

#### Cre line generation

The neuronal (NBT) and oligodendrocyte (Olig1) Cre lines were generated using multisite gateway technology^38^. The neuronal Cre line, Tg(nbt:mtagBFP2-T2A-iCre) was generated as follows. pCAGGS-mTagBFP2-T2A-iCre (v2) (a gift from Mario Capecchi, Addgene plasmid # 125822) was converted into a middle entry gateway clone (pME:mTagBFP2-T2A-iCre) using BP Clonase II (Invitrogen) with primers F 5’ GGGGACAAGTTTGTACAAAAAAGCAGGCTATGGTGTCTAAGGGCGAAGA and R 5’ GGGGACCACTTTGTACAAGAAAGCTGGGTTCAGTCCCCATCCTCGAGCA with plasmid pDONR221. When a correct clone was identified, the pME:mTagBFP2-T2A-iCre was combined with entry vectors (p5E_nbt -Xenopus neural-specific beta tubulin, p3E-polyA (Tol2Kit #302) and destination vector pDestTol2CG2 (Tol2Kit #395) using LR Clonase II (Invitrogen) as previously described^38^. Correct clones were identified by restriction digest and then sequenced to confirm identity.

The oligodendrocyte Cre line Tg(olig1:iCre) was generated in the same fashion using LR Clonase II (Invitrogen) but instead using plasmids p5E_Olig1^39^, pME_iCre (Tol2kit #589), p3E_pA (Tol2Kit #302), and pDestTol2pACrymCherry^40^. Correct clones were again identified by restriction digest and then sequenced to confirm identity.

Transgenic lines were generated by injecting the expression clones at the one cell stage in combination with tol2 transposase mRNA. Individuals expressing BFP in the CNS for Tg(nbt:mtagBFP2-T2A-iCre) or the red eye marker for Tg(olig1:iCre) were selected and raised to adulthood. Founders were identified by crossing each line to WT and identifying those with either blue CNS for Tg(nbt:mtagBFP2-T2A-iCre) or red eye expression Tg (olig1:iCre).

#### MBP Knock In line generation

The GFP P2A MBP knock in line (KI GFP P2A MBP) was generated as previously described^35^ using CrRNA 5’ TTGTCCAGAGGTGCTTGCAG and homology arms 5’ TGTAGACCACTGAACAGATCAACACCTAGA and 5’ GCCACTGCAAGCACCTCTGGACAAAACCCCTTCGGGCTGG. Briefly, GFP P2A was amplified with primers F 5’ TGTAGACCACTGAACAGATCAACACCTAGAATGATGGTGAGCAAGGGCGAGGA GC and R 5’ CAGCCCGAAGGGGTTTTGTCCAGAGGTGCTTGCAGTGGCGGACCGGGGTTTTCTTCCACGTCTCCTGCTTGCTTTAACAGAGAGAAGTTCGTGGCCTTGTACAGCTCGTCCATGCC using X2 Q5 polymerase (NEB). The PCR product was purified via gel and co injected with the guide RNA. GFP positive individuals were identified at 4dpf and raised. Founders were identified by crossing mosaic GFP adults to WT and screening for F1 with GFP positive oligodendrocytes. The latter were sequenced at the MBP locus to confirm correct integration before crossing to other lines for experiments.

#### Crosses and genotyping information

Δnls^Ubi^ crosses were performed as follows TDP-43 +/-;TDPL +/- X TDP-43 Δnls^gene off^/+;TDPL -/+. Each clutch was sorted for RFP expression at 24hpf and then genotyped for TDP-43/TDPL traditional mutations post hoc.

Δnls^neuro^ crosses were performed as follows Tg(nbt:mtagBFP2-T2A-iCre);TDP-43 +/-;TDPL +/- X TDP-43 Δnls^gene on^/+;TDPL -/+. Each clutch was sorted for the BFP hatching pec marker at 24hpf and sorted for RFP expression in the spinal cord post hoc. TDP-43^UFLIP KO; neuro^ crosses were performed as follows Tg(nbt:mtagBFP2-T2A-iCre);TDP-43 +/-;TDPL +/- X TDP-43 UFLIP KO ^gene on^/+;TDPL -/+. Each clutch was sorted for the BFP hatching pec marker at 24hpf and sorted for RFP expression in the spinal cord post hoc. Δnls^oligo^ crosses were performed as follows Tg(olig1:iCre);TDP-43 +/-;TDPL +/- X TDP-43 Δnls^gene on^/+;TDPL -/+. Each clutch was sorted for the BFP hatching pec marker at 24hpf and sorted for RFP expression in the eyes post hoc.

#### Genetic manipulation to prevent myelin formation

To prevent myelination, an olig2 start site morpholino 5’-CGTTCAGTGCGCTCTCAGCTTCTCG-3 (Gene Tools) was injected at the one cell stage as previously described^24^. Tg(mbp:EGFP-CAAX) was always co injected alongside experimental injections, to confirm the morpholino had prevented oligodendrocyte formation and subsequent myelination.

#### Axonal imaging

The Mauthner line Tg(hspGFF62A:Gal4) was utilized for axonal analysis^22^. Axon diameter was assessed using confocal imaging and super-resolution techniques, focusing on the axon above the cloaca as described previously^41,42^. Briefly, larvae were anesthetized with 600 µM tricaine in E3 embryo medium and immobilized in 1.3–1.5% low melting-point agarose on a glass coverslip. This was adhered to a microscope slide using high vacuum silicone grease to create a well with E3 embryo medium and 600 µM tricaine. Confocal z-stacks with optimal z-steps were captured using a Zeiss LSM880 microscope with Airyscan FAST in super-resolution mode, employing a 20× objective lens (Zeiss Plan-Apochromat 20× dry, NA = 0.8), and processed with the default Airyscan settings (Zen Black 2.3, Zeiss). Axonal swelling area was calculated by measuring total area of the swelling 50µm from the tip using Fiji.

### Automated live imaging

The KI GFP-P2A-MBP and Tg(mbp:EGFP-CAAX) were used to quantify oligodendrocyte number and myelin levels respectively using a previously established automated imaging pipeline. Briefly, the Large Particle (LP) Sampler and VAST BioImager (Union Biometrica) automates the transfer of zebrafish from 96 wells plates to the BioImager microscope platform, whereupon control is passed onto a spinning disk confocal microscope for image acquisition^43^. Images were taken with an upright Axio Examiner D1 (Carl Zeiss) microscope equipped with a high-speed CSU-X1 spinning disk confocal scanner (Yokogawa, Tokyo, Japan). All zebrafish were imaged from a lateral view in 6 individual images stitched together during analysis. After imaging each larvae was dispensed back to the 96 well plates for post hoc genotyping. Due to the variability of the Tg(mbp:EGFP-CAAX), readings where normalised to the mean of control individuals from each clutch and expressed as a percentage. For myelin quantification, clutches were kept separate and readings normalised to WT controls. The KI GFP-P2A-MBP is a knock in and therefore consistent expression between individuals, oligodendrocyte numbers were quantified in absolute terms.

### Statistical analysis

All experiments were analysed using GraphPad prism software. N numbers are always representative of individual larvae. Each experiment pooled larvae from at least 3 independent clutches. All data are presented as mean ± standard deviation, *p* values and statistical tests are noted in the figure legends. Significance was defined as *p* < 0.05.

## Funding

The Hardingham and Chandran laboratories are supported by the UK Dementia Research Institute (UK DRI-4001, UK DRI-4003 respectively), which receives its funding from UK DRI Ltd, principally funded by the MRC. SC also acknowledges funding from the LifeArc Philanthropic fund. DAL was supported by a Wellcome Trust Senior Research Fellowship (214244/Z/18/Z), a UKRI Frontier Research Grant EP/Z533890/1, and an MS Society Centre of Excellence Award to the University of Edinburgh.

## Notes

### Competing Interest Statement

The authors have declared no competing interest.

